# Mesencephalic projections to the nucleus accumbens shell modulate value updating during probabilistic reversal learning

**DOI:** 10.1101/2024.02.24.581852

**Authors:** Katharina Zühlsdorff, Sammy Piller, Júlia Sala-Bayo, Peter Zhukovsky, Thorsten Lamla, Wiebke Nissen, Moritz von Heimendahl, Serena Deiana, Janet R. Nicholson, Trevor W. Robbins, Johan Alsiö, Jeffrey W. Dalley

## Abstract

Cognitive flexibility, the capacity to adapt behaviour to changes in the environment, is impaired in a range of brain disorders, including substance use disorder and Parkinson’s disease. Putative neural substrates of cognitive flexibility include mesencephalic pathways to the ventral striatum (VS) and dorsomedial striatum (DMS), hypothesised to encode learning signals needed to maximize rewarded outcomes during decision-making. However, it is unclear whether mesencephalic projections to the ventral and dorsal striatum are distinct in their contribution to flexible reward-related learning. Here, rats acquired a two-choice spatial probabilistic reversal learning (PRL) task, reinforced on an 80%:20% basis, that assessed the flexibility of behaviour to repeated reversals of response-outcome contingencies. We report that optogenetic stimulation of projections from the ventral tegmental area (VTA) to the nucleus accumbens shell (NAcbS) in the VS significantly impaired reversal learning when optical stimulation was temporally aligned with negative feedback (i.e., reward omission). Moreover, the exploitation-exploration parameter, *β*, was increased (indicating greater exploitation of information) when this pathway was optogenetically stimulated after a spurious loss (i.e. an incorrect (20%) response at the 80% reinforrced location) compared to after a spurious win (i.e. a correct (20%) response at the 20% reinforced location). VTA → NAcbS stimulation during other phases of the behavioural task was without effect. Optogenetic stimulation of projection neurons from the substantia nigra (SN) to the DMS, aligned either with reward receipt or omission or prior to making a choice, had no effect on reversal learning. These findings are consistent with the notion that enhanced activity in VTA → NAcbS projections leads to maladaptive perseveration as a consequence of an inappropriate bias to exploitation *via* positive reinforcement.

## Introduction

Cognitive flexibility refers to the capacity of individuals to shift behaviour adaptively to optimise rewarded outcomes. Flexible responding requires dynamic updating of value associated with stimuli and/or actions (O’Doherty, 2011) and depends in part on signaling from midbrain dopamine (DA) neurons (Cools et al., 2001, 2007). The ability to switch behaviour adaptively is compromised in a broad range of neurological and neuropsychiatric disorders, including Parkinson’s disease (PD) (Cools et al., 2001), schizophrenia (Leeson et al., 2009), and obsessive-compulsive disorder (Remijnse et al., 2013).

Supporting a role for DA in cognitive flexibility, depletion of DA in the caudate nucleus of monkeys (Clarke et al., 2011) or homologous dorsomedial striatum (DMS) of rats (O’Neill and Brown, 2007) impaired the flexible reversal of previously learnt stimulus-reward associations. In contrast, local DA receptor activation in the ventral striatum impaired reversal-learning (Verharen et al., 2019), while infusing DA receptor antagonists into the nucleus accumbens shell (NAcS) or core (NacC) facilitated reversal learning (Sala-Bayo et al., 2020). Such findings support the hypothesis that DA neurotransmission in the DMS and nucleus accumbens (NAc) exerts opposing effects on reversal learning. Such divergence may underlie the impairments in probabilistic reversal learning (PRL) produced by L-DOPA and other DA medications in PD patients (Cools et al., 2001; Frank et al., 2004). Conceptually, this may result from the overstimulation of DA receptors in the ventral striatum, which in relation to the dorsal striatum is less affected by DA loss during the early stages of PD (Morrish et al., 1996).

Midbrain DA projections from the ventral tegmental area (VTA) and substantia nigra pars compacta (SNc) to the striatum support the coding of both positive and negative reward prediction errors (RPEs), specifically, the signalling of discrepancies between expected and received rewards and thereby value-based learning of stimuli associated with actions (Chang et al., 2016; Schultz, 1998; Steinberg et al., 2013). Consistent with this idea, chemogenetic activation of the VTA → NAc pathway impaired spatial reversal learning in rats, an effect linked to learning from reward omissions (loss of rewards) (Verharen et al., 2018). However, few studies have investigated how RPEs causally affect associative learning in the context of reversal learning, despite a growing number of studies reporting temporally-aligned neuronal signaling and stimulus-reward learning (Aquili, 2014; Chang et al., 2015; Steinberg et al., 2013).

Here, we sought to demonstrate a causal link between midbrain neuronal activity and reversal-learning performance in rats by dissociating the effects of positive and negative feedback during PRL. Reward on this task was delivered on 80% of correct trials and 20% of incorrect trials (see **Fig.1**). Thus, to maximize food reward, animals were required to discount spurious negative and positive feedback. We used *in-vivo* optogenetics to investigate the effects of activating either the VTA → NAcS pathway or the medial SNc → to DMS pathway on reversal learning performance. Specifically, we determined the impact on reversal learning of pathway-specific activation after each of the following events: (1) the delivery of reward following a correct response; (2) the omission of reward following a correct response; (3) the delivery of reward following an incorrect response; (4) the omission of reward following an incorrect response. We also investigated the effects on reversal learning of pathway-specific activation immediately before the selection of a response (i.e., after the presentation of the response levers).

**Figure 1.**
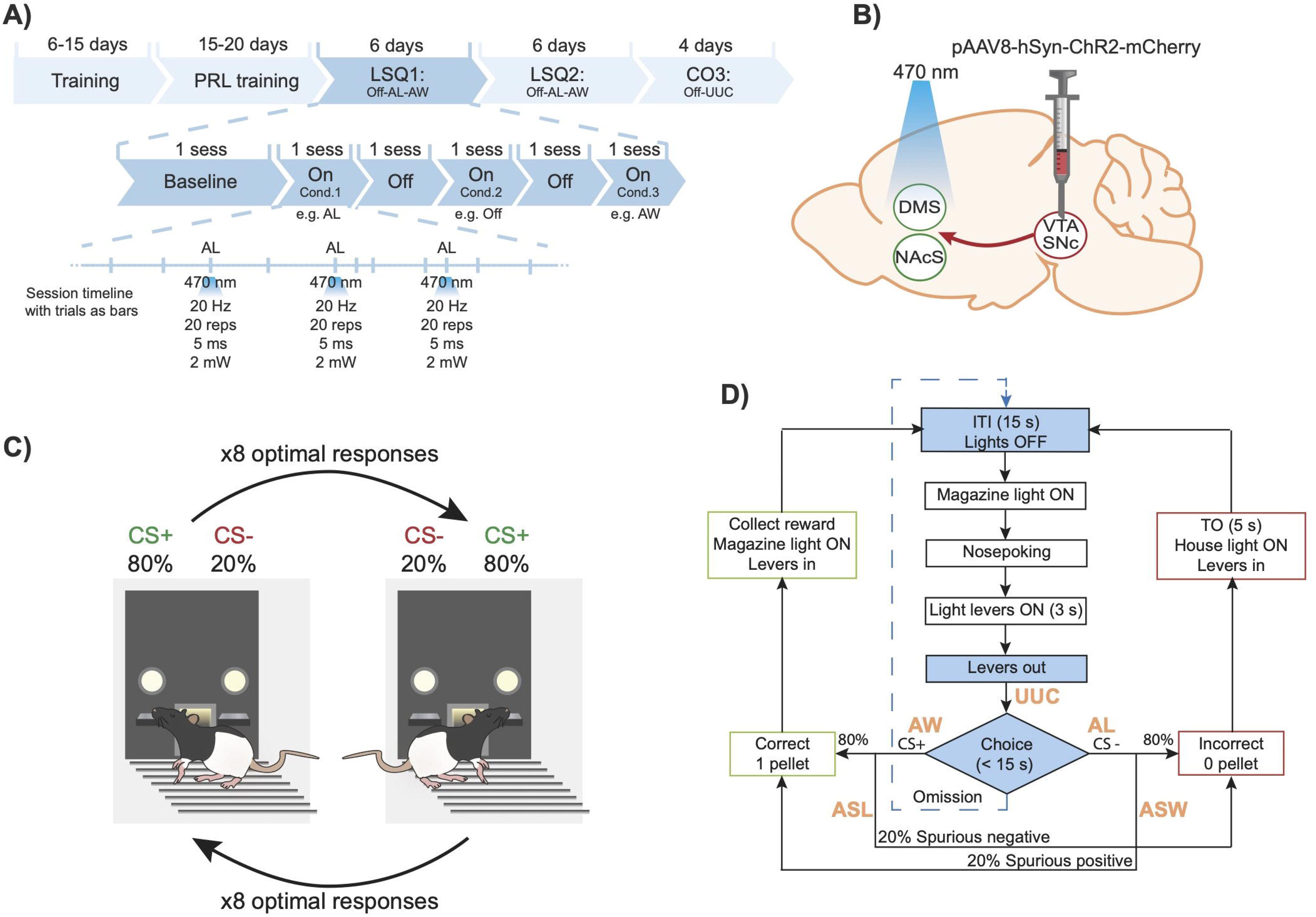
Summary of the experimental procedures used for *in-vivo* optogenetic stimulation of the mesoaccumbal and nigrostriatal pathways during a spatial PRL task. A) Timeline of the behavioural procedure and optical stimulation. Following pre-training stages to respond to the levers, animals were trained in the PRL task. In later sessions of this stage, animals were connected to the patch-cord cables without light stimulation. Once performance had stablilized (i.e. ≥ 3 reversals in 3 sessions in a row), testing started. Testing consisted on two Latin-square experiments and one crossover experiment (LSQ; CO). Each LSQ started with a baseline session consisting on a PRL session with the cables connected to the rats, without light stimulation. The different conditions (e.g. Off-AL-AW) were then tested. Light (*λ* = 470 nm) was selectively turned on after each trial that presented the tested condition e.g. after not receiving a reward (AL). Intracranial stimulation was induced with 20 repetitions of 5-ms pulses, with 2 mW at the tip of the optical fibres. Testing conditions within each LSQ or CO experiment were pseudo-randomized. Final group sizes: DMS (LSQ1) n = 24, (LSQ2) n = 24, (CO3) n = 24; and NAcS (LSQ1) n = 15, (LSQ2) n = 13, (CO3) n = 11. Abbreviations: after loss (AL), after win (AW), after a spurious loss (ASL), after a spurious win (ASW), up until making a choice (UUC), session (sess). B) Schematics showing viral vector infusion in the VTA/SNc, and fibre optic implantation in the NAcS or DMS. C) Illustrative overview of the PRL task in lever-pressing chambers. After eight consecutive correct trials on the optimal lever (CS+; 80% rewarded), the contingencies were reversed such that the previously optimal lever was now suboptimal (CS-; 20% rewarded), and *vice versa*. D) Flowchart of the PRL task. Shown in orange are the time-points where neuronal pathways were optogenetically stimulated for the following events: AL, AW, ASL, ASW, and UUC.

We hypothesised that increased VTA-NAcS neuronal activation timed to coincide with reward omission (i.e., negative RPE) would impair reversal learning by interfering with the putative endogenous dip in DA release that encodes the teaching signal to omitted rewards (Schultz et al., 1997). We predicted that this intervention would decrease shifting behaviour and the rate of learning from unrewarded or negative feedback trials. We further predicted that increasing VTA → NAcS neuronal activation on rewarded trials, thereby mimicking positive RPEs, would either enhance reversal performance by amplifying the presumed natural burst of DA release or would remain unaltered due to possible ceiling effects. Given the opponent effects of dorsal *versus* ventral DA signaling on reversal learning, discussed above, we anticipated that SNc → DMS neuronal activation would result in broadly opposite effects on reversal learning compared with VTA → NAcS pathway activation.

## Materials and methods

### Subjects

Forty-six male Lister-Hooded rats (Charles River, Germany) were initially housed in groups of four under humidity- and temperature-controlled conditions and a 12/12-h light-dark cycle (lights off at 07:30 h). Rats had a minimum of 7 days to acclimatise to the animal facility before any experimental procedures began. Rats were ∼300 g at the beginning of training and were maintained at 90% of their free-feeding weight by food restriction (19 g/day of Purina chow). Water was provided *ad libitum*. Experiments were conducted in accordance with the German animal welfare legislation, Association for Assessment and Accreditation of Laboratory Animal Care (AAALAC) regulations and approved by the Local Animal Care and Use Committee.

### Stereotaxic surgery

Anaesthesia was induced with 5% isoflurane in oxygen and maintained at 2.5%. Rats were secured in a stereotaxic frame fitted with atraumatic ear bars (KOPF Model 1900, Germany). An incision was made along the midline of the skin overlying the dorsal skull. The skull surface was manually cleaned, and OptiBond^®^ All-in-One bone glue (Kerr, USA) was applied and hardened with UV light for 60 s. Animals were bilaterally infused with a maximal total volume of 2400 nL of viral vector, divided across four infusions at a flow rate of 200 nL/min. Recombinant AAVs (rAAV) were produced by transient transfection of adherently grown HEK-293H cells in CELLdiscs, and purified by PEG-precipitation, iodixanol density gradient ultracentrifugation, Amicon-15 ultrafiltration and sterile filtration, as previously described in detail (Strobel et al., 2015, 2019). Genomic titres were determined by qPCR using primers specific to the human synapsin promoter sequence. Optogenetic animals received opsin-expressing rAAV8-hSyn-ChR2-mCherry (7.3 × 10^12^ particles/ml) whereas control animals received rAAV8-hSyn-mCherry (8.38 × 10^12^ particles/ml). Animals were divided into two groups: (1) VTA → NAcS (n=20); (2) SNc → DMS (n=26). For the first group, the virus was infused into the VTA at anteroposterior (AP) -5.4 and -6.2, mediolateral (ML) ± 0.6, dorsoventral (DV) -8.4 and -7.8 and optical fibers (Doric Lenses, Canada) were implanted in the NAcS at AP + 1.5, ML ± 0.8, DV – 7.0. For the second group, the virus was infused into the SNc at AP -5.4, ML ± 0.6; DV -8.1 and -8.0 and optical fibres (Doric Lenses, Canada) were implanted in the DMS at AP + 1.2, ML ± 2.0, DV – 5.3 (**Fig. 1B**). Coordinates refer to millimeters (mm) from Bregma and the skull surface. Infusion cannulae were left in place for 5 min after each infusion to allow for diffusion. Implants were secured with dental cement, four skull screws, and dental product Charisma^®^ (Kulzer, Germany), and hardened with UV light for 20 s to increase the gripping surface for the cement. All surgeries took place at least four weeks before the start of behavioural testing to ensure adequate opsin transfection. After surgery, rats were single housed for the first three days of recovery. They were then pair-housed for the rest of the study. Behavioural training commenced 7 days after surgery.

### Behavioural apparatus

Rats were trained in eight operant chambers (Med Associates, Georgia, VT, USA), each enclosed within a sound-attenuating wooden box fitted with a fan for ventilation and two response levers. Each chamber measured 31.4 x 25.4 x 26.7 cm with a Plexiglas ceiling, front door, and a back panel. On one side of the chamber, a food magazine was centrally placed and equipped with light and a photocell nose-poke detector (**Fig. 1C**). A pellet dispenser was connected to the magazine to deliver the reward: 45 mg sucrose pellets (5TUL, TestDiet, USA). Two retractable levers and a cue light above each lever flanked the magazine. On the same wall, a house light (3W) was positioned. The opposite wall had a metal panel. The floor was made of stainless-steel bars separated 1 cm from each other with a tray underneath. Access was through a hinged sidewall, secured with a latch during testing. Optogenetics cables were connected from the rats to the ceiling of the boxes through a central hole in the ceiling of the chamber (diameter: 5 cm).

### Behavioural training

Training started at least two days after food restriction. Animals were first habituated to the chambers for 15 min with three sugar pellets placed in the magazine before the start of the session. In all stages, trials began with the illumination of the magazine light. Rats were then trained in stage 1 or ‘conditioning’, which consisted of a session of 60 minutes or 40 trials, whichever came first, to learn that pressing the lever delivered reward pellets. Both levers were presented simultaneously, and when one lever was pressed, three pellets were delivered and both levers were withdrawn; if no lever was pressed within 30 seconds, one single pellet was delivered and both levers were withdrawn. The criterion to move to the next stage was the completion of all 40 trials. The following training stages consisted of sessions of 60 min or 120 trials, whichever came first, and incorporated an inter-trial interval (ITI) of 10 s, a time-out (TO) of 5 s after an incorrect response, and a limited hold (LH) of 30 s, after which levers were retracted and the trial was deemed an omission (no free pellets during this or any later sessions). The criterion to move to subsequent stages was the attainment of ≥ 80 correct trials. In stage 2, animals were trained to press the lever to receive a reward (one pellet when one of the two levers was pressed), or none if not. After 60 trials, the preferred lever was retracted to force animals to press the opposite lever and not to develop a side bias. The following stage, stage 3, was similar to the previous stage except that animals had to nose poke into an empty magazine to initiate each trial. The objectives of the final training stage (stage 4) were to avoid side bias and for the rat to learn that both levers were rewarded in a probabilistic manner. For this, both cue lights were illuminated for 3 s, after which only one of the levers was extended, which was rewarded on 80% of the trials. After 30 consecutive trials, the opposite lever was extended for the next 30 trials, and so on until the rat had completed 120 trials. At least two sessions reaching criterion (≥80 correct trials) were required to move to probabilistic reversal learning (PRL) task training.

### Probabilistic reversal learning task

Behavioural training in the PRL task was modified from Bari et al. (2010) for the use of retractable levers instead of nose poking holes (**Fig. 1**). Briefly, daily sessions consisted of 200 trials or 60 min, whichever came first, including an ITI of 10 s, TO of 5 s and LH of 10 s. At the start of each session, one of the two levers was randomly selected to be the ’optimal’ lever, for which responding was more likely to be reinforced. Sessions began with two free pellets in the magazine and illumination of the magazine light. After nose poking, the magazine light was extinguished, and the two cue lights turned on, indicating that the levers would be available three seconds later. A response to the optimal lever delivered a single reward pellet on 80% of trials, whereas a response to the sub-optimal lever yielded reward on only 20% of trials. A failure to press any lever within the LH led to the retraction of the levers and termination of the trial, which was noted as an omission. After eight consecutive correct trials (i.e. pressing the ’optimal’ lever regardless of it being reinforced or not), the contingencies were reversed, so that the previous optimal lever was now sub-optimal and *vice versa* (**Fig. 1C**). This pattern was repeated over the session. Animals were trained until they could achieve at least three reversals per session over three consecutive sessions. Once this criterion was met, rats underwent testing.

### Behavioural testing

Before testing, rats received a minimum of two habituation training sessions with the cables attached to their implant. A baseline session always occurred on the day before testing with the cables attached but with no optical stimulation. For each session, the optimal lever was set as the lever in which rats had finished the previous session to avoid forcing a reversal. Rats were randomly assigned to an optogenetic stimulation group or a ‘light off’ group (**Fig. 1D**). The optogenetic stimulation conditions were: 1) “after loss” (AL) when pressing the sub-optimal lever; 2) “after win” (AW), when pressing the optimal lever; 3) “after spurious loss” (ASL) when pressing the optimal lever but not receiving the expected reward (20% of the times); 4) “after a spurious win” (ASW) when pressing the sub-optimal lever but receiving an unexpected reward (20%); and 5) “up until choice” (UUC) from the start of the trial (presentation of the levers) until pressing a lever (**Fig. 1D**). All conditions were pseudo-randomized according to baseline levels of performance using a Latin-square design (LSQ; (1) Off – AL - AW; (2) Off – ASL – ASW), or cross-over design (CO; (3) Off – UUC) (**Fig. 1A**). Testing took place every second day, leaving 48 h in between each optogenetic session. On intervening days, animals were run with cables attached but with the light off to maintain stable levels of performance and to avoid possible carry-over effects of light stimulation.

### Optical stimulation

Mono fibre-optic patch-cord cables (Doric Lenses, Canada) were metal shielded and terminated in an optical fiber of 200 µm diameter, and a numerical aperture of 0.37. One end of the cable was connected to a PlexBright dual LED commutator *via* magnetic Blue PlexBright compact LED modules (*λ* = 465 nm, max current 200 mA; Plexon, Dallas TX, USA). A computer running Med PC IV (Med Associates) software, which also recorded responses at both levers and magazine, controlled the optical stimulation. A second computer controlled the behavioural task *via* transistor-transistor logic (TTL) signals.

Patch-cord cables were covered with 16 cm plastic tubes to prevent animals from bending and interfering with the cables. Cables were secured to the rats’ implants with a zirconia sleeve (Doric Lenses, Canada) for a 1.25 mm diameter ferrule. Intracranial stimulation was achieved with 20 repetitions of 5-ms light pulses (20 Hz), delivering 2 mW at the tip of each optical fiber (**Fig. 1A**). Data from sessions where light output was compromised because of broken or disconnected optical cables were discarded. Final group sizes were: DMS (LSQ1) n = 24, (LSQ2) n = 24, (CO3) n = 24; NAcS (LSQ1) n = 15, (LSQ2) n = 13, (CO3) n = 11.

### Histological assessment of fibre-optic probe placement and viral vector expression

Following completion of the behavioural procedures, animals were anaesthetized with a lethal dose of pentobarbital (Narcoren, Boehringer Ingelheim GmbH, Germany) and perfused transcardially with 0.01 M phosphate-buffers saline (PBS) followed by 4% paraformaldehyde (PFA) administered with a pump flow rate of 8 ml/min. Brains were removed and post-fixed in 4% PFA for 24 h and dehydrated for cryoprotection in 30% sucrose in 0.01 M PBS.

Brains were coronally sectioned at 60 μm using a cryostat (Leica, Germany), collected in PBS containing 25% polyethylene glycol and 25% glycerol, and stored at 4°C. Free-floating sections were washed in PBS and subsequently blocked and permeabilized in PBS containing 3% normal goat serum (NGS) and 0.3% Triton for 1 h. Sections were incubated overnight with primary antibodies in PBS containing 3% NGS and 0.3% Triton. Since the infused viral vector inherently expressed fluorescence, no antibodies were required to detect transgene expression. For the first set of slices, tyrosine-hydroxylase (TH) was detected with the primary antibody anti-TH in rabbit (1:600, EMD Millipore - Merck, USA). After washing in PBS, sections were incubated with secondary antibodies for 2 h (anti-rabbit in goat Alexa-Fluor 488 nm, 1:500, Invitrogen Thermo Fisher Scientific, USA). For the second set of slices, double staining was achieved with the primary antibodies anti-GAD67 in mouse (1:600, Invitrogen Thermo Fisher Scientific, USA) and anti-VGLUT2 in rabbit (1:600, Invitrogen Thermo Fisher Scientific, USA). Secondary antibodies were goat anti-mouse (Alexa-Fluor 647 nm, 1:500, Invitrogen Thermo Fisher Scientific, USA) and goat anti-rabbit (Alexa-Fluor 488 nm, 1:500, Invitrogen Thermo Fisher Scientific, USA). After washing in PBS, sections were mounted in distilled water and covered with mounting medium (DAPI, EMD Millipore, USA) and a coverslip. Immunofluorescence sections were checked and digitized using a PerkinElmer Opera Phenix High-Contrast Screening microscope (PerkinElmer, USA).

### Computational analysis of behavioural data

To model latent behavioural processes underlying reversal learning performance, we implemented reinforcement learning (RL) algorithms (Zhukovsky et al., 2019), including three variants of the Q-learning model (Daw, 2009) defined below, using three parameters: *α*, *β*, and *κ*. The learning rate *α* determines the degree to which animals learn in response to feedback. The learning rate was further split into *α_win,_* and *α_loss_*. The *α_win_* learning rate determines how quickly the model adjusts the expected Q value of a response following the receipt of a reward (positive feedback), while the *α_loss_* learning rate determines how quickly the expected Q value was adjusted following a non-rewarded response. Expected Q values were converted into action probabilities using the softmax rule by incorporating the inverse temperature parameter *β* and the choice autocorrelation or side “stickiness” parameter *κ*. In this implementation of the model, low *β* values result in random exploration of both response options, and down weigh the contributions of the expected Q values to the probability of choosing a given action. High *β* values result in greater exploitation of the Q values. In the present reversal task, with 80/20 probabilistic outcomes, a low *β* value would result in fewer rewards overall. Finally, the choice autocorrelation parameter *κ* is a measure of side “stickiness”, or how likely it is that an animal will perform the same response again regardless of outcome. Values of *κ* > 0 reflect an agent “sticking” to the previous response while *κ* < 0 reflects choice alternation. In the 80/20 probabilistic reversal task, a moderately high “stickiness” is advantageous as it leads agents to ignore the spurious wins and losses.

#### Model-free Q-learning: Model 1

Simple Q-learning is equivalent to Rescorla-Wagner learning (Rescorla & Wagner, 1972), whereby an agent assigns an expected Q value to each choice available; in the PRL task, a left or right response (L or R) for each trial *t*. The expected Q value for each lever is updated on each trial according to the following:

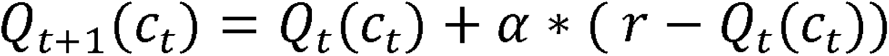

Where 0□≤□α≤□1 is a learning rate parameter, *Q_t_(c_t_)* is the value of the choice *c_t_* at trial *t* and *r* takes the value of *1* if the choice was rewarded and a value of 0 if not. The probability of making the choice *c_t_* at trial *t* was calculated using the softmax rule:

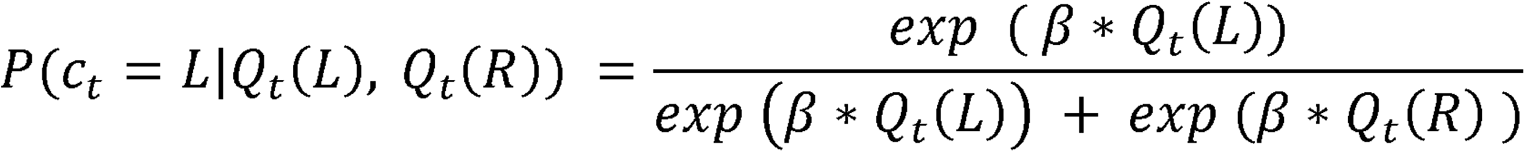

#### Model-free Q-learning: Model 2

Model 2 differed from model 1 only in including the side “stickiness” parameter (*κ*) in addition to *β*:

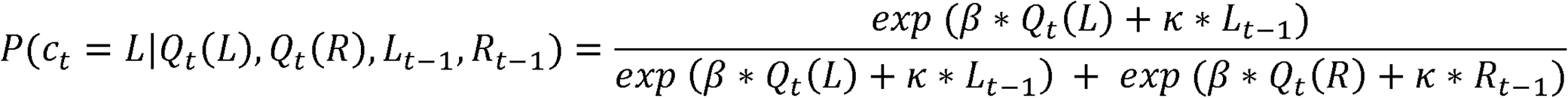

where *L*_t-1_ takes the value 1 if the agent responded left on the previous trial (and otherwise 0), and *R*_t-1_ takes the value 1 if the agent responded right on the previous trial (and otherwise 0). Thus, a larger *κ* results in greater probability of the choice *c_t_* at trial t being the same as the choice *c_t_* at trial *t−1*.

#### Model-free Q-learning: Model 3

Model 2 was extended to include a separate *α* for learning from rewards and losses, *α_win_* and *α_loss_*, depending on whether the animal received a reward or not on trial *t.* The decision probability was updated in the same way as in Model 2.

#### Model fitting and comparison

More details on the model fitting and comparison can be found in (Zhukovsky et al., 2019) and (Daw, 2009). Briefly, the parameters were fitted to maximize the probability of data *D* (the product of the individual probabilities of making a choice *c_t_* at trial *t*) by finding the maximum of the probability density function 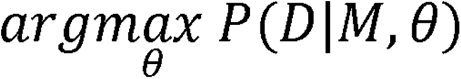.

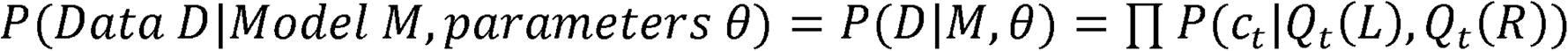

Model space was treated as discrete, using the following range of parameters: 0 ≤ (*α_R_* and *α_NoR_*) □≤1 with a step size of 0.05; 0.005□≤□β□≤□5 with a step size of 0.05 and −1□≤□*κ*□≤□1 with a step size of 0.05. The parameter range was chosen based on *a priori* expectations regarding *α* and *κ*, as well as empirical information about best-fit *β* parameters.

Model selection was conducted using the Bayesian Information Criterion (BIC) that incorporates the likelihood of data given the model with the best fit parameters 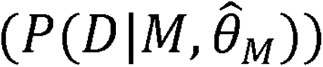 and a penalty term 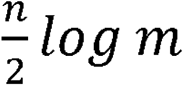:

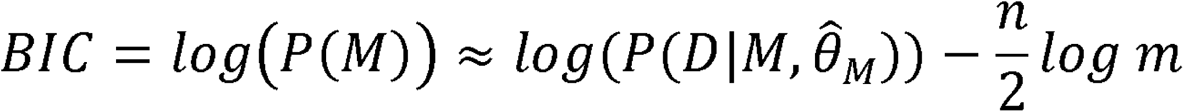

where *n*□=□number of free parameters and *m*□=□number of observations. We also report a biased measure of model fit, *pseudo r^2^*. Scripts implementing the models were written in MATLAB^®^ R2019a and can be found here: https://github.com/peterzhukovsky/reversal_learning. The best model was selected based on the lowest BIC and *pseudo r^2^* values.

#### Model validation using simulations

To further assess the validity of the winning model, we used a set of simulations. Specifically, we simulated groups of rats with mean parameter values selected randomly from each opsin group and light condition in the actual experiment. The number of simulated rats matched the number of actual rats in the particular experiment. Then, each simulated rat completed the PRL task in a virtual environment, updating the Q values and the probabilities of choosing left or right depending on the four individual parameters of that particular group (i.e., *α_win_*, *α_loss_*, *β* and *κ* for winning model 3) and a trial-by-trial accumulation of information, including reward probabilities (i.e., 80%/20%) and reversals after 8 consecutive responses to the optimal lever.

### Behavioural data analysis

The following behavioural measures were extracted and analysed: trials to criterion (TTC), number of reversals, proportion correct responses, percentage of lose-shift behaviour after a correct or incorrect response, percentage of win-stay after a correct or incorrect response, and the average number of perseverative responses after a reversal. The four parameters from the winning Q-learning model were also analysed.

Statistical tests were performed using R version 4.0.4 (R Core Team, 2021). Data were subjected to Linear Mixed-Effects Model analysis with the lmer package in R (Bates et al., 2015). The model contained two fixed factors (experiment – e.g., after win/after loss, light – on/off) and one factor (subject) modeled as an intercept to account for individual differences between rats. Normality was checked with both quantile-quantile (QQ) plots and the Shapiro test. Latencies were log transformed to ensure normality. Homogeneity of variance was verified using Levene’s test. Outliers were identified using Tukey’s method, which highlights outliers ranged above and below 1.5* the interquartile range using the *outlierKD* function in R (Klodian, 2017). The identified outliers were removed from subsequent analyses. When significant interactions were found, further analyses were carried out using *post-hoc* pairwise comparisons using the *emmeans* package in R (Lenth et al., 2018). Significance was considered at p<0.05.

## Results

### Histological analysis

Fig. 2A, B show that administration of opsin-expressing virus into the VTA or SNc resulted in high expression of ChR2 in both the NAcS and the DMS, respectively. Fig. 2C shows fibre optical tip placements in the NAcS or DMS for those animals that completed the study.

**Figure 2.**
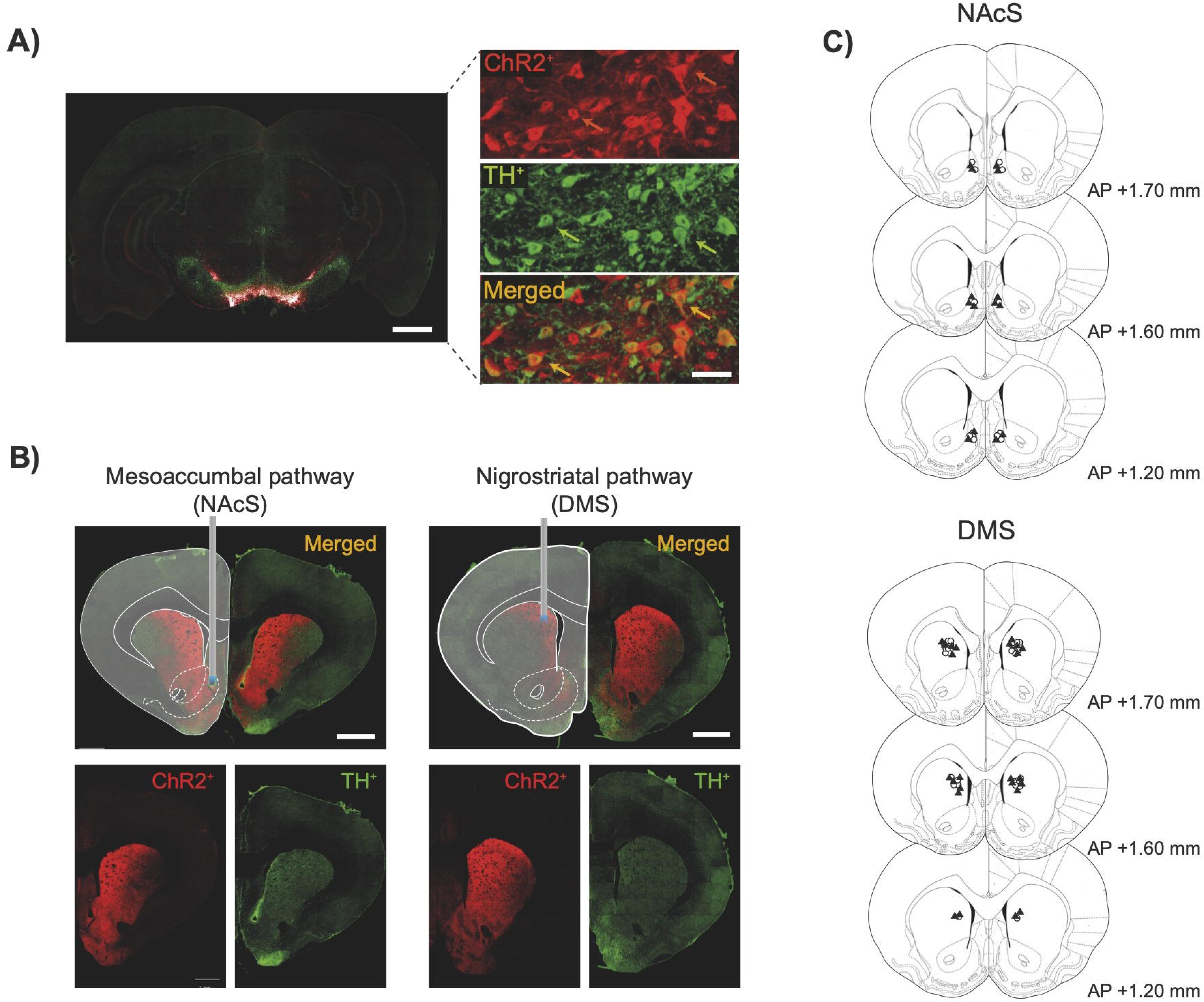
Histological analysis showing viral expression and fibre optic placements in the nucleus accumbens shell and dorsomedial striatum. A) (Left) Coronal section of the ventral tegmental area (VTA) stained for tyrosine hydroxylase (TH) showing viral-transfected neurons. Scale bar: 1 mm. (Right top) Detailed expression of virus positive neurons, (right middle) TH^+^ neurons, and (right bottom) both channels merged. Scale bar: 50 μm. B) Representative histology images showing coronal sections of the striatum and representative fibre optical tip location in the NAcS (left) and DMS (right). Expression of viral vector (ChR2; bottom left), TH (bottom right), and both channels merged (top). Scale bars: 1 mm. C) Fibre optic tip placements in the NAcS and DMS. Full triangles: opsin group. Empty circles: control group. Anteroposterior (AP) coordinates from Bregma.

The expression of virus and neuronal markers in the neuronal fibres was measured in the DMS and NAcS to quantify the neurochemical phenotype of infected neurons. 55.28 ± 3.74% of transfected neurons in the NAcS were TH positive, 15.24 ± 1.56%. were VGLUT2 positive, and 4.77 ± 1.13% were GAD67 positive. In the DMS, 73.51 ± 2.26% were TH positive, 8.99 ± 0.69% were VGLUT2 positive, and 6.17 ± 0.83% were GAD67 positive.

### Behavioural results

#### Activation of the VTA-NAcS pathway impairs reversal learning after reward omission

Optogenetic stimulation of the VTA-NAcS pathway was delivered at different timepoints during the PRL task. With respect to the AL/AW experiment, a near-significant interaction between opsin group and stimulation condition was observed on the number of trials required to reach criterion following stimulation of the VTA-NAcS pathway (group x condition interaction: F_2,42_ =3.07, p = 0.057, η_p_^2^=0.08). Since the p-value for the interaction was less than 0.1, *post-hoc* analyses were carried out according to the guidance of Midway and colleagues (Midway et al., 2020). *Post-hoc* analyses revealed that optogenetic stimulation of the VTA-NAcS pathway selectively impaired performance by increasing the number of trials required to reach criterion compared with the no-light condition when optogenetic stimulation was presented after the loss of reward (t_36_=-3.05, p = 0.045; **Fig. 3**) but not after reward delivery. Moreover, there was a significant group x condition interaction on win-stay behaviour after a correct response (F_2,45_=3.35, p=0.044, η_p_^2^=0.13). None of the *post hoc* analyses were significant following correction for multiple comparisons. However, this intervention had no effect on the proportion of correct responses, win-stay behaviour, lose-shift behaviour or perseverative responses when applied after spurious reward losses or spurious reward wins compared with the light-off condition (i.e., Off *vs* ASL or Off *vs* ASW) or when applied before a response was selected (i.e., Off *vs* UUC).

**Figure 3.**
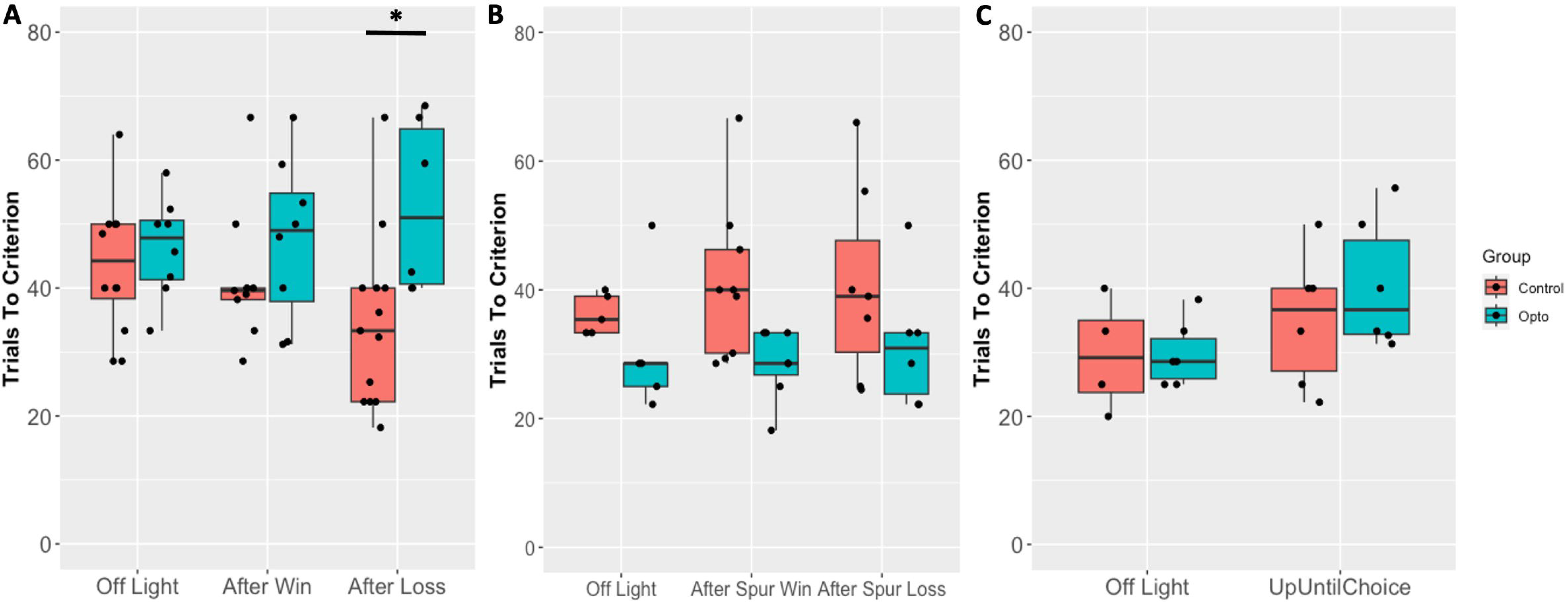
*In-vivo* optogenetic stimulation of the mesoaccumbal pathway impairs reversal learning when selectively activated after the loss of reward. Number of reversals achieved during each session in each experiment: after win/after loss (A), after spurious win/after spurious loss (B), up until choice (C). * - p < 0.05.

With respect to the AL/AW experiment, a significant group x condition interaction was observed with respect to response latencies (F_2,42_=3.52, p=0.039, η_p_^2^=0.11). Response latencies were slower when optostimulation was applied after a win compared to after a loss in the opsin group (t_42_=-2.71, p=0.044). In the other two experiments (ASL/ASW and UUC), there were no effects of condition or group on either the response or collection latencies.

#### Activation of SNc-DMS does not affect reversal learning performance

In contrast to activation of the VTA-NAcS pathway, activation of the SNc-DMS pathway had no significant effects on the number of trials required to reach criterion (**Fig. 4**), number of reversals, lose-shift, number of perseverative responses or any other measure in any of the experiments (AW/AL, ASW/ASL, UUC). There were also no significant group x condition interactions for the response or collection latencies for any of the experiments.

**Figure 4.**
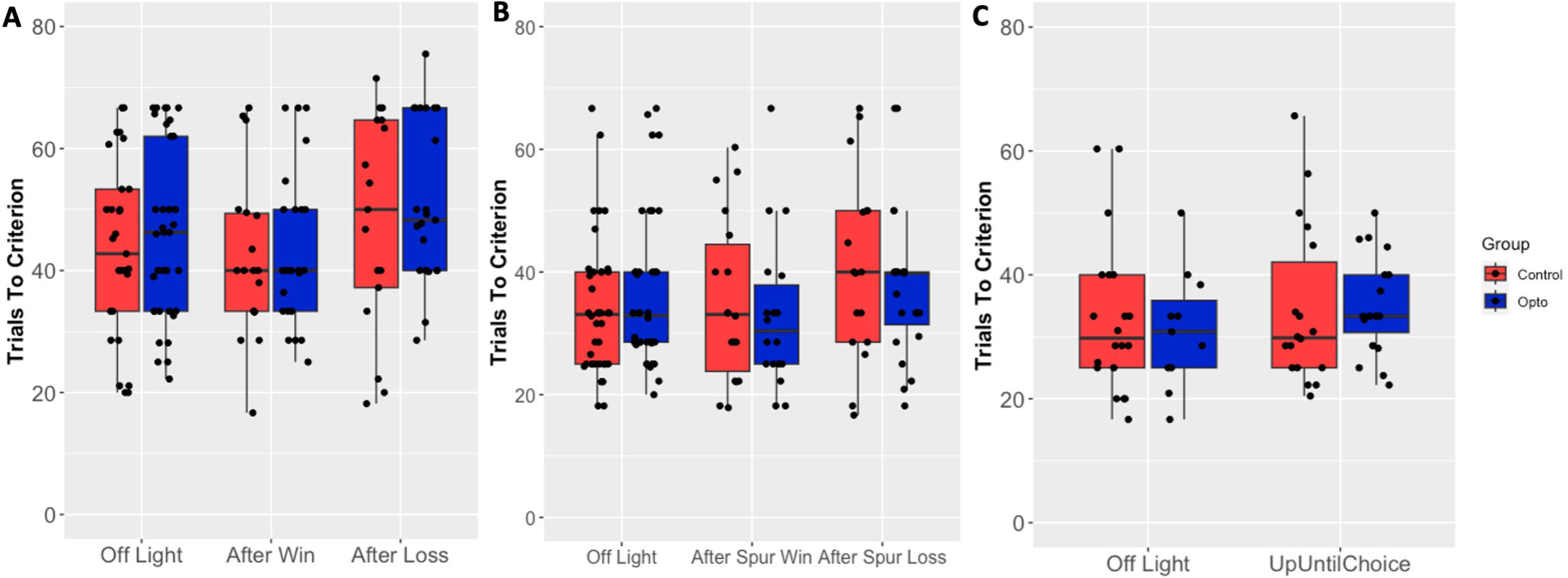
*In-vivo* optogenetic stimulation of the nigrostriatal pathway has no effect on reversal learning performance. Number of reversals achieved during each session in each experiment: after win/after loss (A), after spurious win/after spurious loss (B), up until choice (C). * - p < 0.05.

#### Computational modeling

To explore latent variables influencing behaviour in the spatial PRL task, we used three different reinforced learning algorithms. Model 3 provided the best fits compared to the other two models, since is had lower BIC and *pseudo r^2^* values (Table 1). Model 3 included four free parameters, *α_win_*, *α_loss_*, *β*, and *κ*, which were fitted to the data from each session of each animal, allowing for within-subject comparisons of model performance of sessions with optogenetic stimulation against the light off sessions.

**Table 1.** Summary of model comparison measures for each experiment and each model. Percentage of sessions on which the difference in the Bayesian Information Criterion (BIC) and pseudo r^2^ for the two compared models was below zero (i.e., if the BIC difference is below than zero when models 2 and 1 are compared, model 2 has a lower BIC value and is thus the better fitting model). Model 1 - *α*, *β*; Model 2 - *α*, *β*, *κ*; Model 3 - *α_win_*, *α_loss_*, *β*, *κ*. VTA – ventral tegmental area; NAcS – nucleus accumbens shell; SNc – substantia nigra pars compacta; DMS – dorsomedial striatum.

| Model comparison | VTA-NAcS |  | SNc-DMS |  |
| --- | --- | --- | --- | --- |
|  | *BIC* | *pseudo r<sup>2</sup>* | *BIC* | *pseudo r<sup>2</sup>* |
| 3 – 1 | 67.6% | 99.7% | 59.5% | 99.5% |
| 3 – 2 | 99.1% | 91.5% | 99.0% | 92.7% |
| 2 – 1 | 47.9% | 98.2% | 40.8% | 99.0% |

#### VTA-NAcS pathway stimulation impairs reinforcement sensitivity following a spurious loss

**Fig. 5** reports individual modelled parameters for control and ChR2-expressing rats in the VTA-NAcS experiment for optogenetic stimulation after a spurious win or after a spurious loss. In this experiment, a significant condition x group interaction (F_2,92_=3.22, p=0.045) was revealed with respect to the β parameter. *Post hoc* comparisons revealed that in the opsin group, this parameter was increased when stimulated after a spurious loss compared to a spurious win (t_88_=-3.39, p=0.013), indicating that there is greater exploitation of Q values when optogenetic stimulation occurred after a spurious loss, rather than a spurious win. Prior to correction for multiple comparisons, it was found that the group that received optogenetic stimulation after a spurious loss showed a trend increase *β* values compared to the off light condition in the opsin group (t_90_=-1.92, p=0.058), whereas optogenetic stimulation after a spurious win resulted in lower *β* values compared to the off light condition (t_88_=2.02, p=0.047). There were no significant differences in the parameters following stimulation after a loss, win or when the stimulation was prior to the animal making a choice.

**Figure 5.**
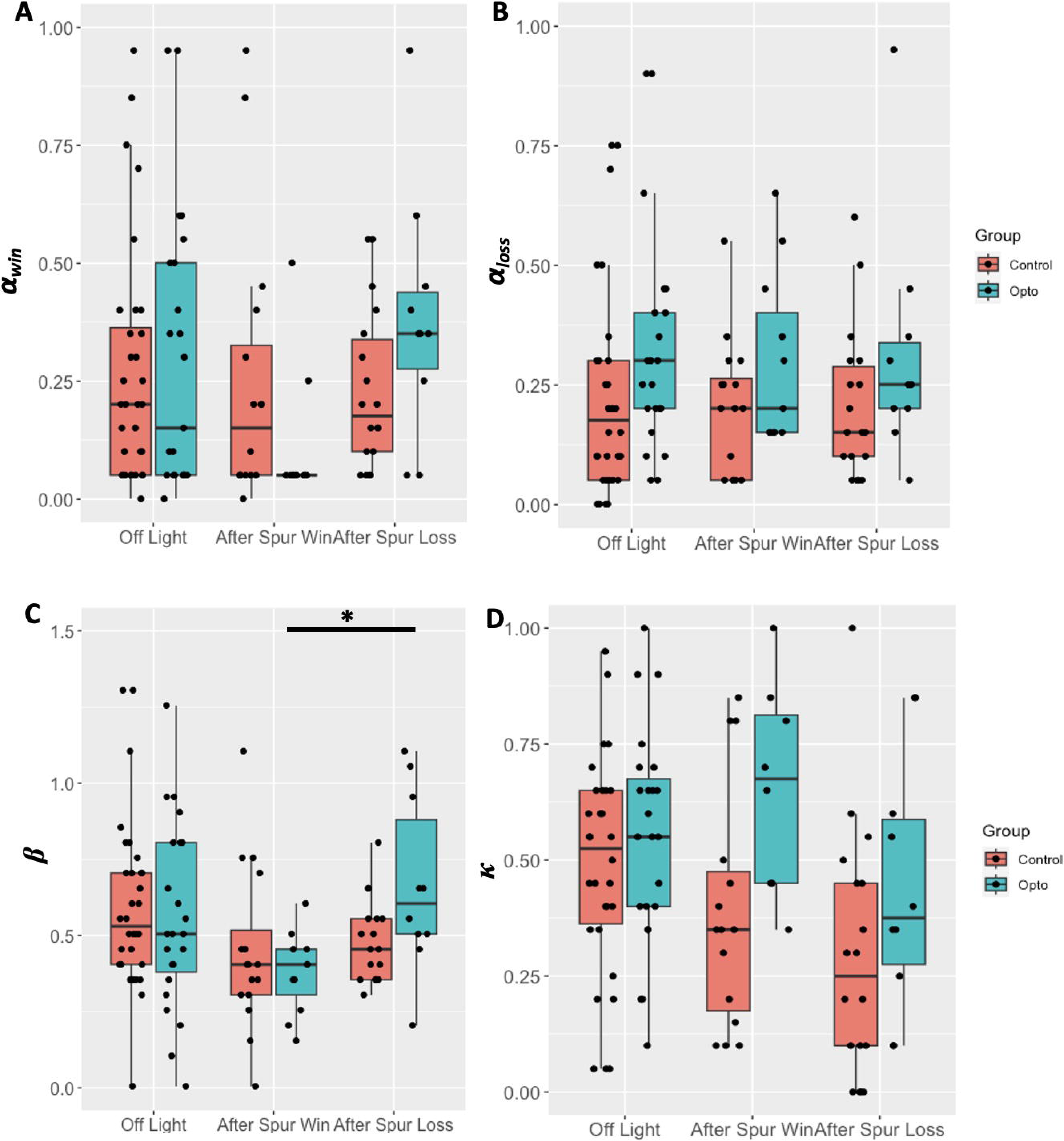
Optogenetic stimulation of the mesoaccumbal pathway after a spurious loss impairs the learning rate from reward absence. When the NAcs was optically stimulated after a spurious win/spurious loss, A) learning rate from rewards (*α_win_*) remained unaltered; B) learning rate from reward absence (*α_loss_*) remained unaltered; C) exploitation/exploration (*β*) parameter was greater in the after spurious loss compared to the after spurious win condition; D) stickiness (*κ*) parameter remained unaltered. * - p < 0.05.

**Figure 6.**
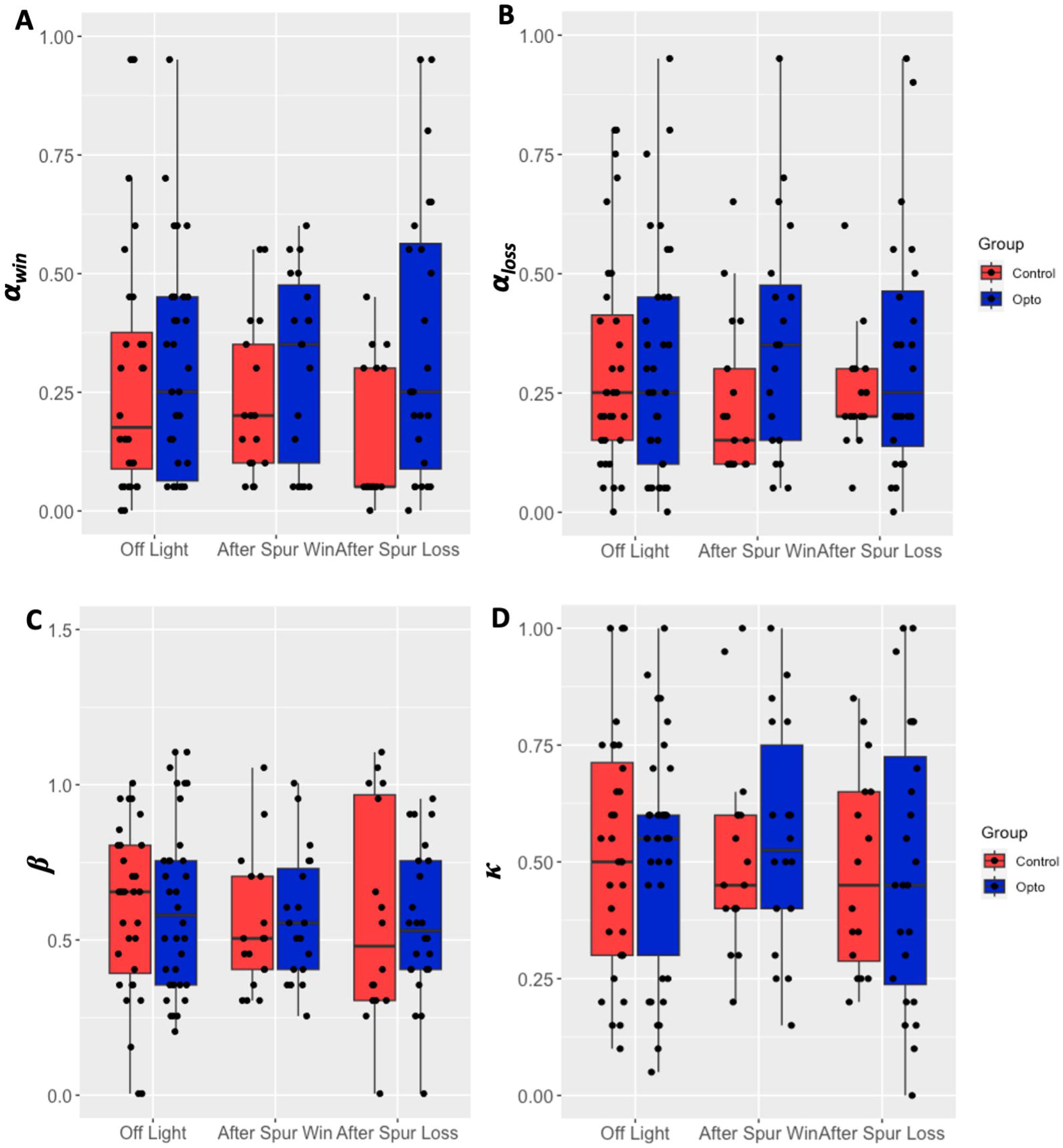
Optogenetic stimulation of the nigrostriatal pathway does not affect reinforcement learning parameters on the PRL task. When the DMS was optically stimulated after a spurious win/spurious loss, A) learning rate from rewards (*α_win_*) remained unaltered; B) learning rate from reward absence (*α_loss_*) remained unaltered; C) exploitation/exploration (*β*) parameter remained unaltered; D) stickiness (*κ*) parameter remained unaltered. * - p < 0.05.

To validate the winning model, we simulated the choice behaviour of agents on the PRL task based on the extracted parameters from model 3. Simulations matched the raw data in the AW/AL experiment, win-stay behaviour after a correct response showed a significant effect of group (F_1,76_=4.34, p=0.041) and condition (F_1,76_=11.29, p<0.001). Moreover, lose-shift after an incorrect response showed a significant group x condition effect (F_2,78_=3.47, p=0.036). No other significant effects were found in the other experiments.

#### SNc-DMS pathway stimulation does not affect reinforcement learning parameters

We found no significant intreactions of main effects on any of the parameters from the winning model (model 3) following optogenetic stimulation of the SNc-DMS pathway after a win or a loss, after spurious win or spurious loss, or up until a choice was made. Simulations matched the behavioural data: there were no significant group x condition interactive effects on any of the conventional measures.

In summary, optogenetic activation of the VTA-NAcS pathway after a loss of reward significantly impaired reversal learning performance by affecting the number of trials required to reach criterion. Moreover, there was an increase in the exploitation-exploration parameter, as indicated by an increase in the *β* parameter, when the VTA-NAcS pathway was stimulated after a spurious loss compared to a spurious win. In contrast, stimulation of the SNc-DMS pathway had no significant effects on any of the behavioural or computational variables.

## Discussion

Using *in-vivo* optogenetic stimulation we report, for the first time, dissociable roles of the VTA-NAcS and SNc-DMS pathways in modulating the effects on reward feedback in a two-choice spatial discrimination reversal task with probabilistic reinforcement. Whereas stimulation of the VTA-NAcS pathway following reward omission (i.e., after loss of reward) impaired reversal learning performance by increasing the number of trials required to reach criterion, no such changes were observed after stimulation of the SNc-DMS pathway. These findings are consistent with the hypothesis that hyperactivation of the VTA-NAcS pathway impairs reversal learning specifically due to aberrant signals following reward omission, and that the NAcS and the DMS have dissociable roles in modulating reversal learning.

The dissociable effects of nigrostriatal DA stimulation are consistent with the NAcS and the DMS having dissociable but complementary roles in modulating reversal learning depending on learning phase (Sala-Bayo et al., 2020). The DMS is implicated in goal-directed behaviours, making it necessary to develop and implement strategies to solve the task. In addition, a study employing a stimulus-response task identified the dorsal striatum as a key region involved in the initiation and execution of a learned instrumental response rather than the encoding of RPEs (Cox & Witten, 2019). Instead, the utilization of RPEs appear to be selectively processed by the NAcS, as suggested by the impairment related to negative feedback processing in the present study.

Using a complementary RL modelling approach, we further found that optogenetic stimulation of VTA-NAcS neurons after a spurious loss resulted in increased exploitation of feedback information as shown through an increase in the *β* parameter. This impairment may be explained by increased baseline levels of DA, caused by optogenetic activation, leading to an interference of negative RPEs carried by dips in DA release following unexpected reward omission (Schultz et al., 2017). This interpretation parsimoniously extends the present findings where temporally precise optogenetic stimulations were used to drive midbrain input to the NAcS during behavioural events (negative RPEs) previously associated with sudden dips in DA release in the NAcS. The presumed lack of a neurochemical signal for negative RPEs would thus be expected to impair the value updating and subsequent suppression of actions directed toward the non-rewarded stimulus, thereby reducing the number of reversals. Such effects are also observed in PD patients following the administration of the indirect DA agonist L-DOPA (Cools et al., 2007).

While we observed that hyperactivation in the NAcS after reward omission impaired performance, an improvement in behaviour as measured through exploitation of Q values was also found after optogenetically activating the pathway after spurious losses. As these trials represent the lack of reward following a response to the optimal lever (i.e., negative RPEs), this aligned with our expectations of animals shifting less when presented with such negative feedback. Unsurprisingly, none of the measures were affected following optogenetic stimulation after an expected or a spurious reward, as the stimulated pathway was already activated by the animal’s experience. One might perhaps have expected hyperactivation to increase the burst of neuronal spiking and thus facilitate reaching the threshold for acting as a teaching signal in positive RPEs, to improve performance. Since no effect was observed, it is reasonable to assume that the system reached ceiling levels of RPE magnitude.

In the present study, a role for non-DA VTA neurons cannot be excluded. Although the virus was mainly expressed in TH^+^ neurons, it was not restricted to these neurons and was present in non-TH^+^, including GAD67^+^ and VGLUT2^+^ neurons. The observed outcomes could thus result from the combined activity of DA, GABA, and glutamate. A convincing body of work has related DA to reinforcement learning, but a recent study suggested a DA-independent contribution by such processes by reporting that glutamate release from glutamate/dopamine co-releasing VTA ➔ NAcS neurons promotes reinforcement in the form of optogenetic self-stimulation (Zell et al., 2020). In addition, DA terminals co-release glutamate preferentially in the ventromedial, but not the dorsal striatum (Mingote et al., 2019; Stuber et al., 2010; Tecuapetla et al., 2010). It is possible that the observed effects after manipulating this region were due to additional gluatamate modulation – perhaps explaining the lack of effect when targeting the DMS. Our results are in line with the improved behavioural flexibility after inhibiting NAcS neurons (Aquili et al., 2014) and possibly the enhanced response switching induced by systemic amphetamine (Evenden & Robbins, 1983). A small proportion of neurons co-expressed the virus and GAD67. Stimulation of VTA-GABA projections to the NAc have been shown to enhance stimulus-outcome learning, likely by inhibiting cholinergic neurons in the striatum (Brown et al., 2012). However, increased stimulus salience would result in improved performance, unlike the observed deficit in reversal learning. Hence, combined with the finding that GABA neurons were in a minority of neurons expressing the virus, it is unlikely the observed deficit in reversal learning was the result of increased GABA signaling in the NAcS.

A further consideration is the potential reinforcing effect of light applied via optogenetics, which could potentially exert rewarding effects. However, under similar conditions, the findings of Steinberg et al (2013) refuted the possibility of conditioned reinforcing effects of the optogenetic stimulation. Additionally, when paired with reward (e.g., in rewarded trials, AW “after win”), the light could increase reward value as a conditioned reinforcer, leading to increased discriminability from the non-rewarded stimulus. However, performance in reversal learning did not improve when either the mesolimbic or nigrostriatal pathways were stimulated during reward delivery; nor was there an increased value or preference for paired rewards (Steinberg et al., 2013). This suggests that optical stimulation was not sufficient to simulate the properties of the natural reward.

The present research expands and supports the role of midbrain-striatal circuit encoding RPEs as teaching signals to enable learning. For the first time, it demonstrates with temporal precision the link between hyperactivity within the VTA-NAcS pathway and performance in reversal learning when negative outcomes are encoded. Our work adds to a burgeoning body of literature establishing and characterizing the causal link between DA signaling during RPEs and reward learning (Aquili, 2014; Chang et al., 2015; Steinberg et al., 2013). Further understanding of the processes involved in learning from negative feedback may be relevant for patients with major depressive disorder, schizophrenia, or OCD, who show an accentuated bias towards negative feedback (Clark et al., 2009; Elliott et al., 1997; Hales et al., 2014).

## Acknowledgements

We thank Ms. N. Okogun and Ms. A. Kaun for skilled technical assistance.

